# Genome-wide strategies reveal target genes of Npas4l associated with cardiovascular development in zebrafish

**DOI:** 10.1101/461988

**Authors:** Michele Marass, Arica Beisaw, Claudia Gerri, Francesca Luzzani, Nana Fukuda, Stefan Günther, Carsten Kuenne, Sven Reischauer, Didier Y. R. Stainier

**Affiliations:** Department of Developmental Genetics, Max Planck Institute for Heart and Lung Research, Bad Nauheim, Germany; ECCPS Bioinformatics and Deep Sequencing Platform, Max Planck Institute for Heart and Lung Research, Bad Nauheim, Germany

**Author notes:** Present address: The Francis Crick Institute, London, UK.

## Abstract

The development of a vascular network is essential to nourish tissues and sustain organ function throughout life. Endothelial cells (ECs) are the building blocks of blood vessels, yet our understanding of EC specification remains incomplete. zebrafish *cloche/npas4l* mutants have been used broadly as an avascular model, but little is known about the molecular mechanisms of action of the Npas4l transcription factor. Here, to identify its direct and indirect target genes, we combined complementary genome-wide approaches including transcriptome analyses and chromatin immunoprecipitation (ChIP). The cross-analysis of these datasets indicates that Npas4l functions as a master regulator by directly inducing a group of transcription factor genes crucial for hematoendothelial specification such as *etv2*, *tal1* and *lmo2*. We also identified new targets of Npas4l and investigated the function of a subset of them using the CRISPR/Cas9 technology. Phenotypic characterization of *tspan18b* mutants reveals a novel player in developmental angiogenesis, confirming the reliability of the datasets generated. Collectively, these data represent a useful resource for future studies aimed to better understand EC fate determination and vascular development.

## INTRODUCTION

The cardiovascular system is one of the first vertebrate organs to develop, and it is critical for the delivery of oxygen and nutrients and the removal of waste products from developing and mature organs (Marcelo et al., 2013). Endothelial cells (ECs) constitute the building blocks of the vasculature, lining the luminal side of all vessels. EC specification is a crucial event for the development of the circulatory system and the lack of a vascular system leads to early embryonic lethality in most animals (Lee et al., 2008). In zebrafish, EC precursors were identified in the ventral region of the gastrulating embryo as early as shield stage (Vogeli et al., 2006). During gastrulation, a subset of ventral mesodermal cells differentiates into angioblasts, the progenitor of ECs. During somitogenesis, the angioblasts migrate from the lateral plate mesoderm (LPM) towards the midline of the embryo, where they coalesce to form the axial vessels: the dorsal aorta and the cardinal vein (Jin et al., 2007). Sequential transcriptional waves orchestrate EC differentiation starting from 6 hours post fertilization (hpf) and a number of transcription factor genes have been described to be involved in this process, including *cloche/npas4l* (Stainier et al., 1995;Reischauer et al., 2016), *fli1* (Craig et al., 2015), *etv2* (Pham et al., 2007), *lmo2* (Weiss et al., 2012), *sox7* (Hermkens et al., 2015) and *tal1* (Bussmann et al., 2007). Importantly, *cloche* mutants exhibit a severe impairment of EC specification, lacking most endothelial and hematopoietic tissues (Stainier et al., 1995), and they fail to activate the EC specification program (Liao et al., 1997;Ho et al., 1999).

The role of *cloche* in EC specification was determined to be cell-autonomous (Parker and Stainier, 1999). For more than two decades, the *cloche* mutant has been employed as an avascular model to study the impact of vasculature on the development of other organs, including the retina (Dhakal et al., 2015), the pancreas (Field et al., 2003a) and the liver (Field et al., 2003b). Since the *cloche* locus is situated in a telomeric region, the isolation of the gene was challenging, and only recently, *cloche* was found to encode a bHLH-PAS transcription factor (Reischauer et al., 2016). The Cloche protein was thereafter called Npas4l due to its partial similarity, in terms of amino acid sequence, to mammalian Npas4 (Reischauer et al., 2016). Interestingly, *npas4l* is present in fish, reptiles and birds, but is missing in mammals, raising questions about its mammalian functional ortholog and the evolution of endothelial cell differentiation (Reischauer et al., 2016).

In this study, we identified genes downstream of Npas4l. We generated novel loss- and gain-of-function models for *npas4l* and combined transcriptomic data with chromatin immunoprecipitation (ChIP) of Npas4l to elucidate its molecular mechanism of action. Our data indicate that Npas4l drives EC differentiation by inducing the expression of the transcription factor genes *etv2*, *tal1* and *lmo2*, directly binding to their promoters. Furthermore, the analyses performed led to the identification of a potential new player in cardiovascular development. Using the CRISPR/Cas9 technology, we generated a mutant for *tspan18b*, which we show is a target of Npas4l. *tspan18b* mutants exhibit early angiogenic defects, confirming the reliability of the datasets generated. Collectively, these data represent a valuable resource of Npas4l target genes that may play a role in vascular and hematopoietic development.

## RESULTS AND DISCUSSION

### Npas4l promotes endothelial cell specification by directly inducing angioblast genes

In order to gain insight into the gene regulatory network controlled by Npas4l *in vivo*, we first conducted an overexpression experiment by injecting *npas4l* mRNA into one-cell stage zebrafish embryos. Following *npas4l* overexpression, we performed microarray analysis to identify transcripts upregulated at 30 and 95% epiboly (**Fig. 1A**). Gene Set Enrichment Analysis (GSEA) results showed that *npas4l* was sufficient to drive the expression of a number of genes involved in blood vessel morphogenesis (**Fig. 1B**), consistent with previous data (Reischauer et al., 2016). As early as 4 hpf (30% epiboly), we detected the induction of angioblast and endothelial markers including *etv2*, *tal1* and *lmo2* (**Table I**). These and other angioblast specific genes were upregulated at both stages (**Fig. 1C**). Additionally, we identified *dnajb5*, a gene encoding a member of the heasthock 40 protein family, which was induced at 30 and 95% epiboly (**Fig. S1A**). *in situ* hybridization analysis showed that *dnajb5* is expressed in the LPM at the 5-somite stage (**Fig. S1B**). Notably, genes expressed in mature endothelium such as *kdrl*, *cdh5, she*, and *clec14a*, which have previously been reported to function downstream of Etv2 (Gomez et al., 2012), were substantially induced at 95% but not at 30% epiboly (**Fig. 1C**).

**Figure 1.**
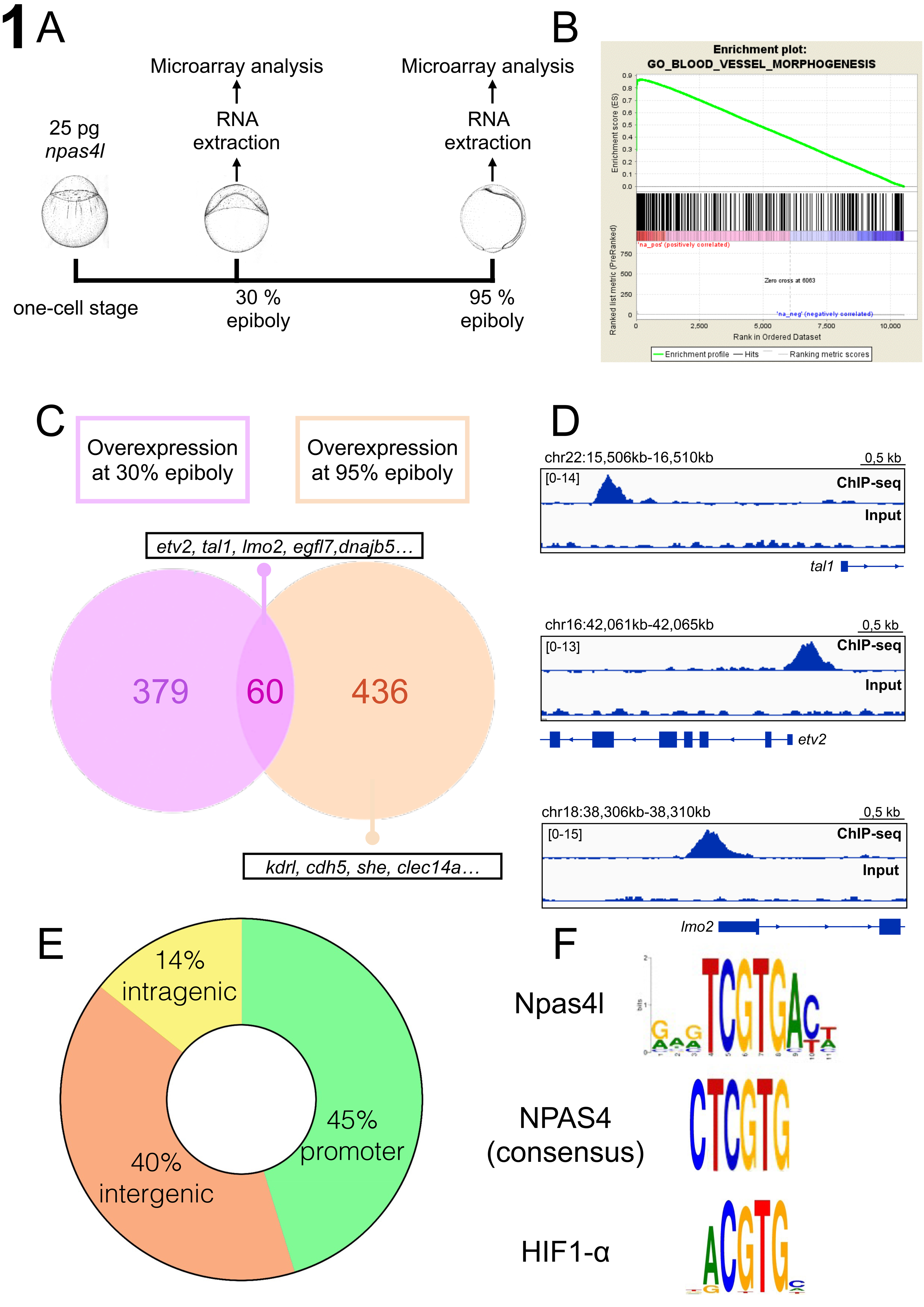
Npas4l promotes endothelial cell specification by directly inducing angioblast specific genes. A)Schematic representation of the overexpression experiment followed by microarray analysis. B) Gene set enrichment analysis (GSEA) performed on upregulated genes at 95% epiboly. C) Venn diagram depicting the genes induced more than 3-fold after *npas4l* overexpression at 30 and 95% epiboly. D) Integrative Genomics Viewer (IGV) visualization of Npas4l occupancy at selected loci: *tal1*, *etv2* and *lmo2*. E) Pie chart showing the proportion of regions bound by Npas4l relative to their genomic localization. F) *de novo* binding motif identification of Npas4l and comparison with previous reported binding motifs of other bHLH-PAS transcription factors.

**Table I.** Overview of the selected genes regulated by Npas4l. List of selected genes across several datasets. Tables are color-coded based on the Venn diagram shown in Figure 3.

|  | *etv2* | *tal1* | *lmo2* | *sox7* | *egfl7* | *cdh5* | *tspan18b* | *si:dkey-1*<br>*21a11.3* |
| --- | --- | --- | --- | --- | --- | --- | --- | --- |
| 30%<br>epiboly<br>uninjected<br>raw value | 2.74 | 2.74 | 7.31 | 10.48 | 2.48 | 6.31 | 47.17 | 2.91 |
| 30%<br>epiboly<br>injected<br>raw value | 2203.40 | 1758.65 | 1613.35 | 399.48 | 45.93 | 14.69 | 50.08 | 11.15 |
| 30%<br>epiboly<br>microarray<br>fold<br>change | 805.05 | 640.72 | 220.80 | 38.11 | 18.50 | 2.33 | 1.06 | 3.83 |
| 95%<br>epiboly<br>uninjected<br>raw value | 18.37 | 7.09 | 21.70 | 61.32 | 2.55 | 2.36 | 154.63 | 26.45 |
| 95%<br>epiboly<br>injected<br>raw value | 14120.60 | 5299.57 | 1937.16 | 2523.17 | 2204.36 | 149.91 | 507.82 | 62.64 |
| 95%<br>epiboly<br>microarray<br>fold<br>change | 768.84 | 747.37 | 89.29 | 41.15 | 863.39 | 63.53 | 3.28 | 2.37 |
| 6 somite-stage<br>WT<br>raw value | 269.00 | 164.50 | 210.50 | 55.00 | 43.00 | 19.50 | 115.00 | 23.50 |
| 6 somite-stage<br>*npas4l*<br>mut<br>raw value | 106.50 | 10.50 | 47.00 | 22.50 | 15.50 | 7.00 | 81.50 | 18.00 |
| 6 somite-stage<br>RNA-seq<br>fold<br>change | 2.53 | 15.67 | 4.48 | 2.44 | 2.77 | 2.79 | 1.41 | 1.31 |
| Tailbud<br>stage<br>Heat<br>shock WT<br>raw value | 190.00 | 266.00 | 190.50 | 103.50 | 46.50 | 131.00 | 174.00 | 34.00 |
| Tailbud<br>stage<br>Heat<br>shock<br>*HOTCre*<br>raw value | 453.00 | 369.50 | 368.50 | 305.50 | 64.50 | 88.00 | 199.50 | 76.00 |
| Tailbud<br>stage<br>RNA-seq<br>fold<br>change | 2.38 | 1.39 | 1.93 | 2.95 | 1.39 | 0.67 | 1.15 | 2.24 |
| ChIP:<br>promoter<br>within 2 kb<br>from<br>TSS | ✓ | ✓ | ✓ |  | ✓ |  | ✓ |  |
| ChIP:<br>promoter<br>within 100<br>kb from<br>TSS | ✓ | ✓ | ✓ |  | ✓ |  | ✓ | ✓ |

The transcriptional analysis performed at two different time points provides insight into the temporal dynamics of the transcriptional cascade initiated by Npas4l. We hypothesized that genes upregulated at 30% epiboly may reflect direct targets, while those upregulated at 95% epiboly represent indirect or secondary targets. In order to determine the direct downstream targets of Npas4l, we performed a genome-wide chromatin immunoprecipitation sequencing analysis (ChIP-Seq). Due to the lack of commercially available antibodies recognizing Npas4l, we used an HA-tagged version of the protein that was able to induce expression of *etv2* at similar levels when compared to wild-type (WT) Npas4l (**Fig. S1C**). We found that Npas4l-HA binds a total of 42 genomic regions, 19 within 5 kb from a transcription start site (TSS) (**Fig. S1D**), 6 in intragenic regions and the remaining 17 in intergenic regions (**Fig. 1E**). Next, we performed a cross-analysis of genes induced at least 2-fold by overexpression of *npas4l* and genes whose TSS is present within 100 kb from an Npas4l binding site, leading to the discovery of 15 genes that are likely to be direct targets of Npas4l, including *etv2, tal1, lmo2, egfl7, gata2a* and *tspan18b* (**Fig. 1D** and **S1E**). We also extrapolated the binding motif of Npas4l using the MEME Suite online tool (http://meme-suite.org), and identified the consensus sequence TCGTGA (**Fig. 1F**). This binding motif is consistent with the non-palindromic binding sites reported for other bHLH-PAS transcription factors, including HIF-1α, characterized by the core sequence CGTG (Schodel et al., 2011;Lo and Matthews, 2012). In particular, the TCGTG site was found to be the target of Dysfusion, the *Drosophila* ortholog of Npas4l (Jiang and Crews, 2007), and the same sequence was described to be the target of the neuronal-specific activity-dependent NPAS4 in mouse (Sorensen et al., 2016). Intriguingly, Tsang et al. (2017)reported that HIF-1α binds to the *Etv2* promoter in human embryonic stem cells and that hypoxia promotes *Etv2* expression via HIF-1α. Considering the similarity of the HIF-1α motif to that of Npas4l (**Fig. 1F**), this bHLH-PAS transcription factor may, at least in part, have taken over *npas4l* function in mammals.

### Transcriptomic and epigenomic analyses of the npas4l mutant

To better describe the molecular mechanisms of Npas4l function, we analyzed the transcriptome and epigenome of *npas4l* mutants. Using CRISPR/Cas9 technology, we generated a new *npas4l* mutant allele with a lesion in its second exon which encodes the bHLH domain (**Fig. S3**). *TgBAC(etv2:EGFP*)^ci1^ *npas4l^−/−^* exhibit a strong phenotype, comparable with the previously described *m39* (Stainier et al., 1995) and *s5* (Liao et al., 2000) alleles, with the advantage that this new allele (*bns297*) can be genotyped using high resolution melt analysis (Samarut et al., 2016). Previous studies had analyzed the transcriptome of *npas4l*/*cloche* mutants at 15- and 18- somite stage (Qian et al., 2005;Sumanas et al., 2005). However, in order to focus on the early expressing genes, we performed RNA-sequencing (RNA-seq) analysis of mutant embryos and their WT siblings at the 6-somite stage (**Fig. 2A)**. *tal1*, *etv2* and *lmo2* were strongly downregulated in the mutant embryos compared to WT siblings (**Fig. 2B**, **Table I**). The expression levels of hematopoietic genes, including *gfi1aa* and *gata1a*, were also decreased in the mutants, although these genes were not induced by *npas4l* overexpression (**Fig. 2C**). Gene set enrichment analysis highlighted a number of genes involved in vascular and hematopoietic development downregulated in *npas4l* mutants at the 6-somite stage, confirming its role in the early specification of endothelium and blood (**Fig. S2A**). Surprisingly, we found that genes involved in neural function were upregulated in *npas4l* mutants, suggesting that angioblasts, or *npas4l* itself, may have a role in repressing neuronal gene expression at 6-somite stage (**Fig. S2B**). Murine NPAS4 is known to regulate synaptic function (Spiegel et al., 2014), and *npas4a* expression was previously described to be restricted to the zebrafish brain, similar to mammalian *Npas4* (Klaric et al., 2014). We previously reported that *npas4l* was expressed specifically in the LPM during early somitogenesis using *in situ* hybridization (ISH) (Reischauer et al., 2016). Nevertheless, neural progenitors might express levels of *npas4l* too low to detect by ISH. Interestingly, a subset of genes regulated by *npas4l*, including *gata2a* and *tal1*, were found to be expressed during zebrafish development not only in EC progenitors, but also in subsets of neurons (Andrzejczuk et al., 2018). Additionally, mesodermal cells may acquire a neural fate in absence of *npas4l* function, similarly to what was reported in the *Tbx6* knockout mouse (Chapman and Papaioannou, 1998).

**Figure 2.**
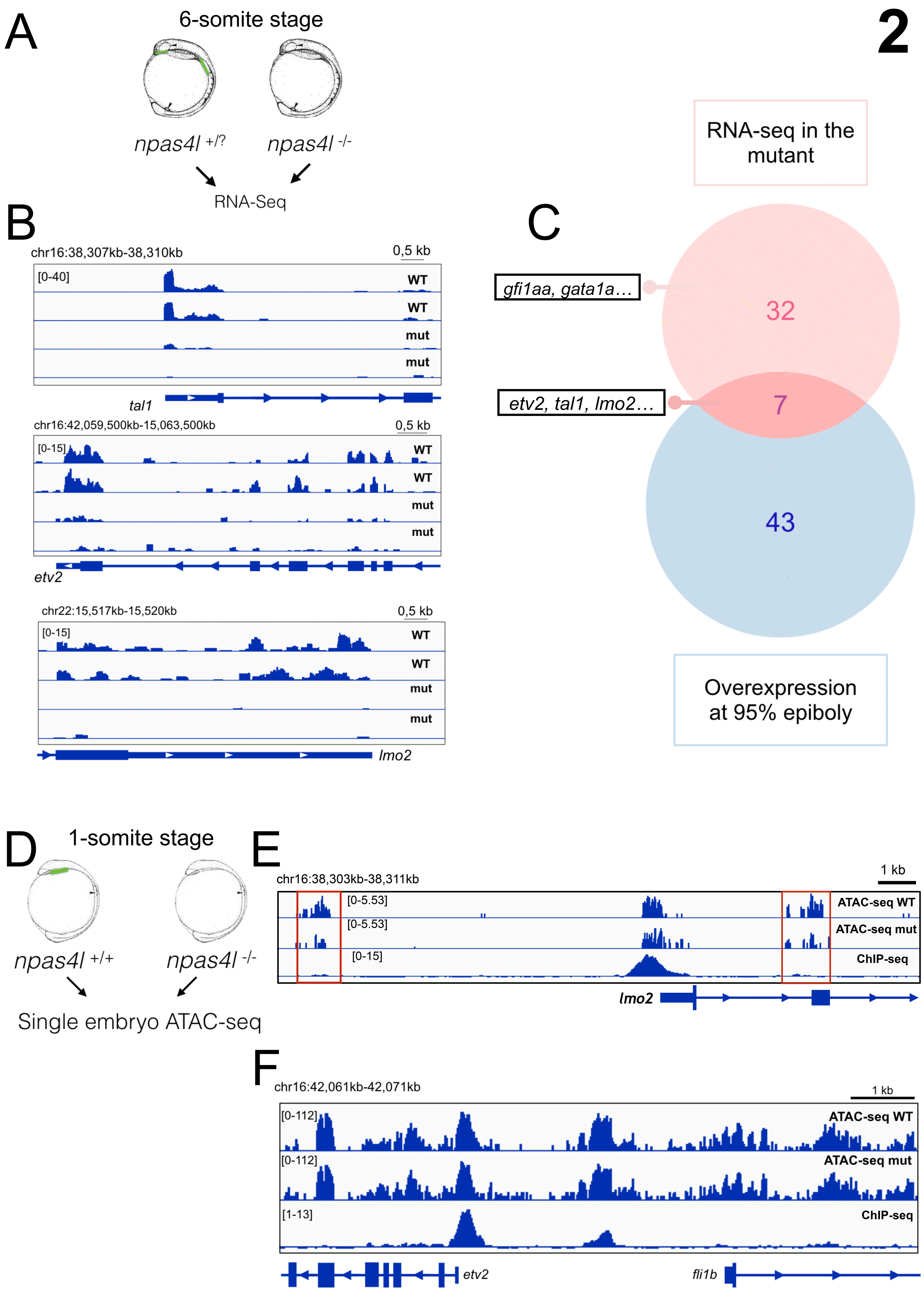
Transcriptomic and epigenomic analyses of *npas4l* mutants. A) Schematic representation of the RNA-seq experiment. B) IGV visualization of selected targets of Npas4l, showing a significant reduction in expression level in *npas4l* mutants. C) Venn diagram showing the genes downregulated in *npas4l* mutants more than 2-fold and the top 50 genes induced by *npas4l* overexpression at 95% epiboly. D) Schematic representation of the ATAC-seq experiment. E) IGV visualization of accessible chromatin at the *lmo2* locus in 1- somite stage WT and mutant embryos, combined with ChIP-seq data. The less accessible regions in mutants are outlined in red. F) IGV visualization of accessible chromatin at the *etv2* locus in 1-somite stage WT and mutant embryos, combined with ChIP-seq data.

Given the striking phenotype of *npas4l* mutants and the ability of Npas4l to potently activate its target genes as early as 4 hpf (**Fig. 1C**), we wanted to test whether Npas4l acts as a pioneer transcription factor. Pioneer transcription factors bind to condensed chromatin and have the ability to make it more accessible, enabling other factors to access their target sites (Iwafuchi-Doi and Zaret, 2014). One possible approach to investigate this question is to assess accessible chromatin in a mutant setting, using the Assay for Transposase-Accessible Chromatin coupled to next generation sequencing (ATAC-seq) (Buenrostro et al., 2015;Doganli et al., 2017). We incrossed *npas4l*^+/−^ fish and genotyped single embryos at the 1-somite stage. Subsequently, we performed ATAC-Seq on mutants and WT siblings (**Fig. 2D**). We compared ATAC-seq data to Npas4l ChIP-Seq results, and observed that the chromatin accessibility at Npas4l binding sites was not affected in *npas4l* mutants compared to WT (**Fig. 2F**). Interestingly, in a small subset of Npas4l targets, chromatin accessibility was reduced, as in the case of *lmo2*, although the affected regions do not appear to interact directly with Npas4l (**Fig. 2E**). The present data indicate that Npas4l binds to open chromatin to recruit the transcriptional machinery, promoting the expression of its target genes. Importantly, Npas4l might regulate chromatin state specifically in angioblasts, and this effect might be diluted in the whole embryo analysis.

### npas4l overexpression at tailbud stage promotes the expression of etv2 but not of cdh5

To test for genes that may be induced by *npas4l* at later stages, we generated and used a heatshock inducible line (**Fig. S4A**). Considering the physiological expression pattern of *etv2*, *tal1* and *lmo2*, we induced *npas4l* expression at tailbud (TB) stage, concomitantly with the appearance of the first angioblast markers (**Fig. S4B**). We performed qPCR at different time points to identify the ideal time point when direct targets were induced, but secondary targets were not affected. At 1 hour post heatshock (hph), *etv2* upregulation was detectable while *cdh5* expression, a known target of *etv2*, was not significantly upregulated (**Fig. S4C**). We performed RNA-seq analysis at 1 hph, and could detect *etv2* expression increasing by more than 2-fold (**Table I**). Notably, *tal1* and *lmo2* expression levels were also upregulated after heatshock, but due to their high level of expression in WT, the induction of these genes was below 2-fold (**Table I**).

### *tspan18b* as a novel gene potentially involved in vascular development

Following the generation and analysis of the individual datasets, we combined the obtained data in order to investigate the mechanisms underlying *npas4l* function and identify new *npas4l* target genes. The resulting Venn diagram includes the overexpression experiment, the transcriptome of *npas4l* mutants at the 6-somite stage and the ChIP-Seq data, considering all genes whose TSS is found within a 100kb window from at least one Npas4l binding site. As the Venn diagram highlights (**Fig. 3A**), using a threshold of 2-fold change, *etv2* is the only gene consistently present in all datasets. Notably, *lmo2*, *tal1* and *egfl7*, despite being direct targets of Npas4l, are not induced by *npas4l* overexpression at TB stage, due to their physiologically high expression level at the stage of sample collection. According to our ChIP-Seq data, Npas4l does not seem to bind to the promoter of *sox7*, although *sox7* is present in all other datasets and it was previously described to act downstream of *etv2* (Wong et al., 2009). Our analysis aimed to identify novel genes and intriguingly, a previously undescribed gene, *si:dkey-121a11.3*, was detected in most of the datasets, although Npas4l does not appear to bind its proximal promoter (**Table I**). We therefore focused our attention on *tspan18b*, a gene induced by *npas4l* and with an Npas4l binding site in its promoter (**Table I**). *tspan18* belongs to the tetraspanin family and it has been reported to be expressed in the developing vasculature in chick (Fairchild and Gammill, 2013) and mouse (Scialdone et al., 2016). To study its function in zebrafish, we generated a mutant line using CRISPR/Cas9 technology by targeting exon 4 and recovered a 16-nucleotide insertion that leads to a premature stop codon after a 74 amino acid-long missense segment (**Fig. S5**). When crossed to the a *Tg(fli1a:EGFP)* background (Lawson and Weinstein, 2002), we observed defects in the formation of the intersegmental vessels at 36 hpf in *tspan18* mutants (**Fig. 3B**). Notably, the phenotype observed was partially penetrant (**Fig. 3C**), possibly due, at least in part, to compensatory mechanisms described to occur following gene knockout (Rossi et al., 2015).

**Figure 3.**
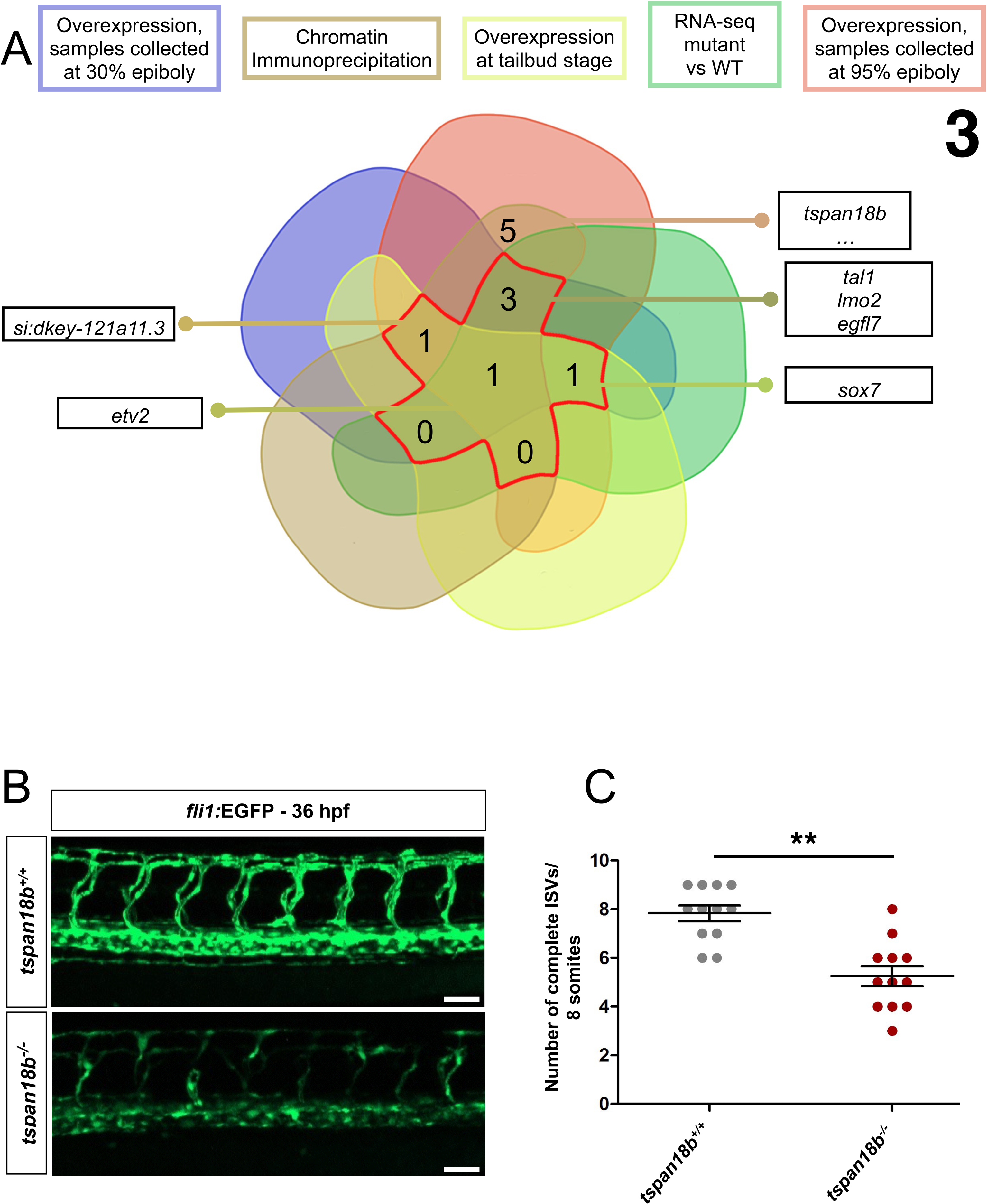
*tspan18b* as a novel gene potentially involved in vascular development. A) Venn diagram illustrating the cross-analysis of the different datasets used to identify genes downstream of Npas4l. B) Maximal intensity projections of confocal Z-stacks of 24 hpf *Tg(fli1a:EGFP)* WT sibling and *tspan18b* mutant embryos. C) Quantification of the intersegmental vessels connected to the dorsal longitudinal anastomotic vessel in *tspan18b*^−/−^ mutants and WT siblings. Each dot represents the number of complete vessels in one embryo. *N* = 24 embryos from three different clutches; (\*\**P*<0.01, *t-test*). Scale bar, 50 μm.

In conclusion, our results represent a collection of genome-wide data that allow for the identification of genes acting downstream of Npas4l *in vivo*. We foresee that the data generated and the analyses performed in the present study will be a valuable resource for the scientific community, especially those who aim to investigate not only EC specification, but also hematopoietic and vascular development. In addition, the present study describes the molecular mechanisms underlying Npas4l function by revealing its binding motif and its direct targets. The analysis of *npas4l* target genes led to the identification and characterization of *tspan18b*, a novel regulator of vascular development in fish, and future studies will be required to further investigate the role of this gene in vascular development and function.

## Acknowledgements

We thank Ryota Matsuoka, Kenny Mattonet, Oliver Stone and Jason Lai for helpful discussions and sharing reagents and protocols.

## Author contributions

M.M. and D.Y.R.S. conceived and directed the study. M.M., S.R., A.B. and D.Y.R.S. designed and supervised experiments. M.M., A.B., C.G., F.L. and N.F. performed experiments. S.G., C.K. and M.M. performed bioinformatics analyses. M.M. and D.Y.R.S. wrote the manuscript. All authors contributed to data analysis and commented on the paper.

